# Early Pitx2 modulation of Fgf10 directs asymmetric lung morphology

**DOI:** 10.64898/2026.06.16.732783

**Authors:** Rui Yan, Jared Helms, Pulin Li, Clifford J. Tabin

**Affiliations:** Department of Genetics, Blavatnik Institute, Harvard Medical School, Boston, MA 02115, USA; Whitehead Institute for Biomedical Research, Cambridge MA, 02142; Department of Biology, Massachusetts Institute of Technology, Cambridge MA, 02139

**Author notes:** Herbert Wertheim School of Public Health and Human Longevity Science, University of California, San Diego, La Jolla, CA 92093, USA. Correspondence should be addressed to R.Y. and C.J.T.

## Abstract

Most of the visceral organs are anatomically asymmetric across the left-right axis. These asymmetries can be traced to a well-studied molecular cascade leading to left-sided expression of a series of genes in the early embryo. One of the key genes in this left-sided cascade is the transcription factor Pitx2, which is transiently expressed throughout the left lateral plate mesoderm. Yet how these early differences in gene expression are converted into differential shaping of organs at later stages remains incompletely understood, and for many organs, such as the lung, the question has not even been explored. Similarly, while the signaling pathways responsible for the branching morphogenesis of the lung have been intensively studied, no insight has been reported regarding how the described gene regulatory network is modified to produce differences in morphology on the left and right sides. Here we identify Fgf10 as a Pitx2-sensitive signal in the mesenchyme of the developing mouse lung. Fgf10 expression increases as Pitx2 decreases, making the right lung, which lacks Pitx2 expression, grow faster than the left during the budding stage. Modulating Fgf10 dosage in the left mesenchyme is sufficient to alter lung budding asymmetry. At the cellular level, Fgf10-dependent lung growth is due to increased levels of epithelial proliferation, without significant differences in directional migration into the mesenchyme. Conditional genetics further show that Pitx2 activity is required exclusively during the early budding stage. Early Pitx2 is sufficient to establish differences in the primary airway branches. These differences in initial conditions are sufficient to direct the subsequent asymmetric branching patterns that eventually manifest as the left-right differences in lung morphology. Thus, Pitx2 converts left-right mesenchymal identity into organ asymmetry by quantitatively tuning Fgf10-dependent epithelial growth during early lung organogenesis.

## Introduction

Although externally, vertebrates are bilaterally symmetric, many of their visceral organs often develop stereotyped left-right asymmetries in size, position, and shape^1^. This asymmetric configuration both serves physiological functions, such as in the specification of the left and right chambers of the heart, and efficiently packs visceral organs inside the body. The mammalian lung is a bilateral organ exhibiting highly conserved left-right asymmetry, with the left lung smaller and having fewer lobes than the right^2^, which is thought to help accommodate the left-positioned heart within the thoracic cavity. As the left and right lung buds arise from the same foregut epithelium but subsequently grow and branch differently, the lung provides a useful system for investigating how robust left-right differences in organ morphogenesis are generated.

Previous work has shown that the homeobox transcription factor PITX2, a conserved effector of NODAL-dependent left-sided patterning, controls the left-right asymmetric morphogenesis of visceral organs across evolution^1^. In the mouse, homozygous knockout of Pitx2 leads to abnormal heart looping, gut rotation defects, and notably, duplication of the right lung on both sides with near-complete penetrance^3–5^. These phenotypes indicate that PITX2 is required for left-sided organ morphogenesis. However, the downstream mechanisms by which PITX2 directs asymmetric morphogenesis differ between organs. In the gut, PITX2 promotes mesenchymal cell condensation in the left mesentery to initiate rotation^6^. In the heart, PITX2 regulates left-right patterning of cardiac structures, including the atria and outflow tract, during asymmetric heart looping^7^. This has not been addressed in the lung. It is known that PITX2 is specifically expressed in the developing left lung mesenchyme^8^. However, how such left-specific mesenchymal gene activity is converted into quantitative differences in the pattern of epithelial growth and branching remains unresolved.

Lung morphogenesis has been extensively studied as a model of branching morphogenesis instructed by reciprocal epithelial-mesenchymal interaction^2,9,10^. In the mouse, lung development starts at around Embryonic Day 9.0 (E9.0) to E9.5 when the left and right lung buds emerge from the ventral side of the foregut, a process dependent on BMP, WNT, and FGF signaling. As the lung buds elongate, they initiate branching, starting with the right lung, which begins to generate primary branches around E11.0, while the left lung remains comparatively simple. Subsequently, the left lung also forms primary branches, followed by continued, stereotypic secondary branching on both sides^11^. Aspects of the mechanisms controlling lung branching are still debated, but many of the key activities have been revealed. For example, the branching process requires biochemical and biophysical interactions between the lung epithelium and mesenchyme, part of which is mediated by mesenchyme-produced FGF10 and epithelium-derived SHH signaling^10–12^. As these are the pathways that control lung morphogenesis, it is reasonable to suppose that their gene network is modulated to generate left-right asymmetric lung growth and branching, but how they connect to PITX2 in the embryonic lung has so far remained elusive.

To that end, we interrogated the morphogenetic mechanisms of lung asymmetry downstream of PITX2. We find that PITX2 limits Fgf10 expression in the left lung mesenchyme, and that the asymmetry in FGF signaling induces differential proliferation, but not directional migration, of the lung epithelium. Intriguingly, PITX2 is only required for lung asymmetry until E10.0, well before branching morphogenesis of the lung begins, suggesting that the asymmetric branching pattern is determined at the early stage of budding, and that these differences in starting conditions of the left and right lung buds are responsible for the subsequent asymmetries in lung morphology. Together, these findings support a model in which PITX2 converts left-right mesenchymal identity into organ-scale asymmetry by tuning Fgf10 dosage and epithelial growth during the initial budding stage.

## Results

### Left-right asymmetry emerges during the initial lung budding stage

To capture the initial symmetry-breaking mechanism in the lung, we performed whole mount immunostaining of the lung epithelium between E9.5-E10.5, which revealed morphological asymmetry at the initial lung budding stage at E9.5 (Figure 1A). We next asked whether the two lung buds differ in morphogenetic forces at this stage. Although the epithelium has been speculated to drive lung bud outgrowth based on the observation that isolated lung epithelium can spontaneously elongate and branch^13,14^, previous work has not evaluated the driving force of lung budding in situ. To probe forces at the epithelium-mesenchyme interface, we performed laser ablation of the sub-epithelial mesenchyme and measured the movement of epithelium after mesenchymal ablation^15^ (Figures 1B and 1C, Video S1). Following mesenchymal ablation, the epithelium moved outward, indicating that the lung bud epithelium exerts an expansive force that is mechanically resisted by the surrounding mesenchyme. Importantly, the total post-ablation displacement of the epithelium was greater on the right than on the left, indicating a left-right difference in the net expansive force (Figure 1D). These results demonstrate that both lung bud morphology and epithelial biomechanics are already asymmetric at E9.5. Since PITX2, the key regulator of lung asymmetry, is exclusively expressed in the lung mesenchyme (Figure 1E), while the loss of PITX2 expression results in a change in the branching morphogenesis of the lung epithelium, these findings suggest that early epithelial asymmetry is likely mediated through epithelial-mesenchymal crosstalk.

**Figure 1.**
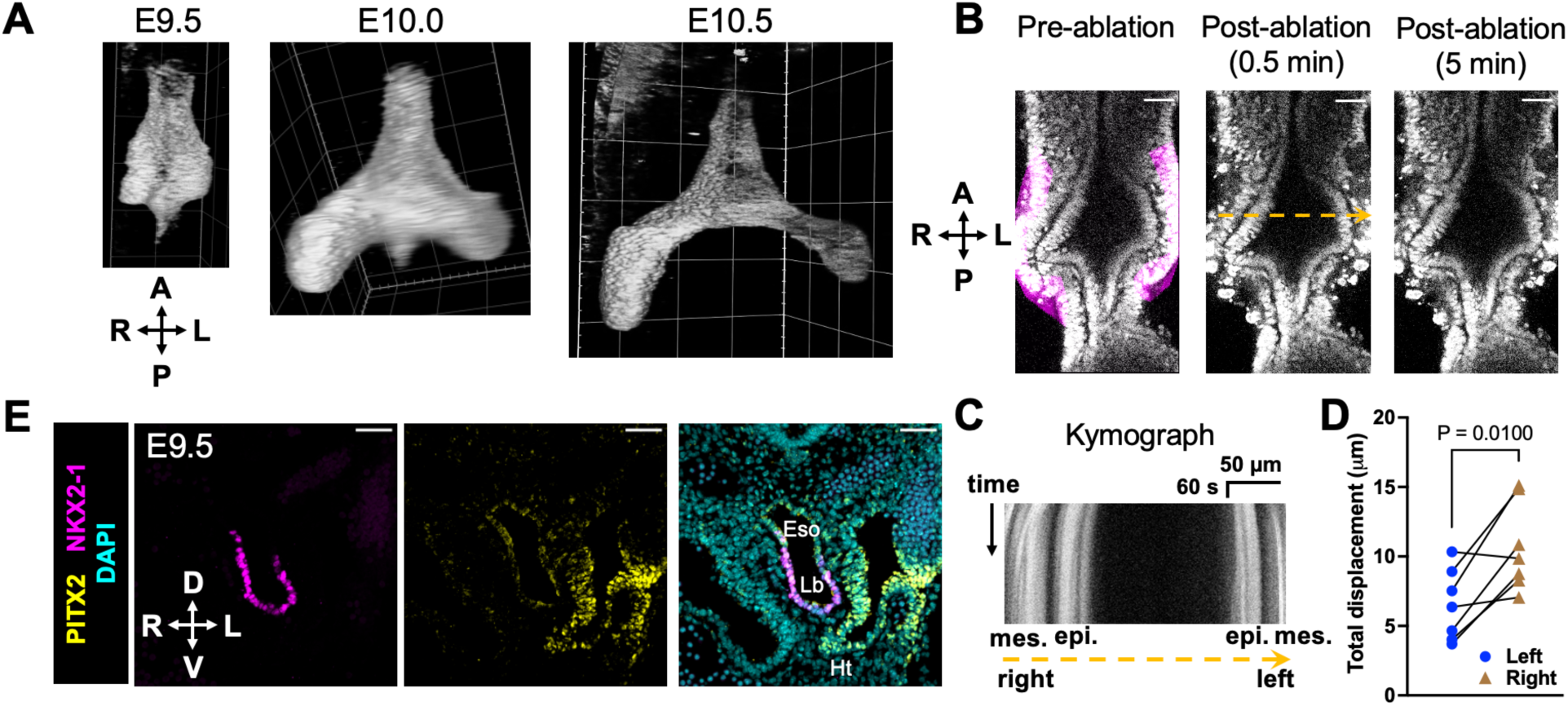
Asymmetric epithelial expansion during lung budding. **(A)** Frontal views of whole mount immunofluorescence of NKX2-1 in early lung buds. Images are representative for N = 5 embryos. Grid size: 100 µm. **(B)** Two-photon laser ablation of left and right sub-epithelial mesenchyme in an E9.5 lung explant. N = 7 embryos. Scale bars: 50 µm. **(C)** Kymograph along the dotted line in (B). **(D)** Quantification of epithelial displacement after ablation. Each data point represents one sample. N = 3 litters. P value: Paired t test. **(E)** Immunofluorescence of NKX2-1 and PITX2 in a transverse section of E9.5 mouse embryo at the lung bud level. Eso: esophagus. Lb: lung buds. Ht: heart. Images are representative for N = 5 embryos. Scale bars: 50 µm. A: anterior. P: posterior. R: right. L: left. D: dorsal. V: ventral.

### PITX2 patterns Fgf10 asymmetry to promote asymmetric lung budding

To identify the asymmetric molecular transducer of PITX2 asymmetry, we sampled the left lung buds from control and Pitx2-knockout (Pitx2-KO) embryos at E10.5, the earliest stage at which the lung bud could be reliably isolated by dissection, and performed bulk RNA sequencing (Figure 2A). We then conducted differential gene expression analysis, which revealed genes altered by PITX2 in the lung bud, many of which have unclear developmental functions (Figure 2B, Table S1). Gene network analysis of the STRING database highlighted FGF signaling-associated genes, including Fgf10, among transcripts increased in Pitx2-KO lung buds (Figure 2C), suggesting FGF signaling as a candidate pathway regulated by PITX2, while BMP, WNT, and SHH pathways did not show significant changes in this dataset (Figures S1A and S1B).

**Figure 2.**
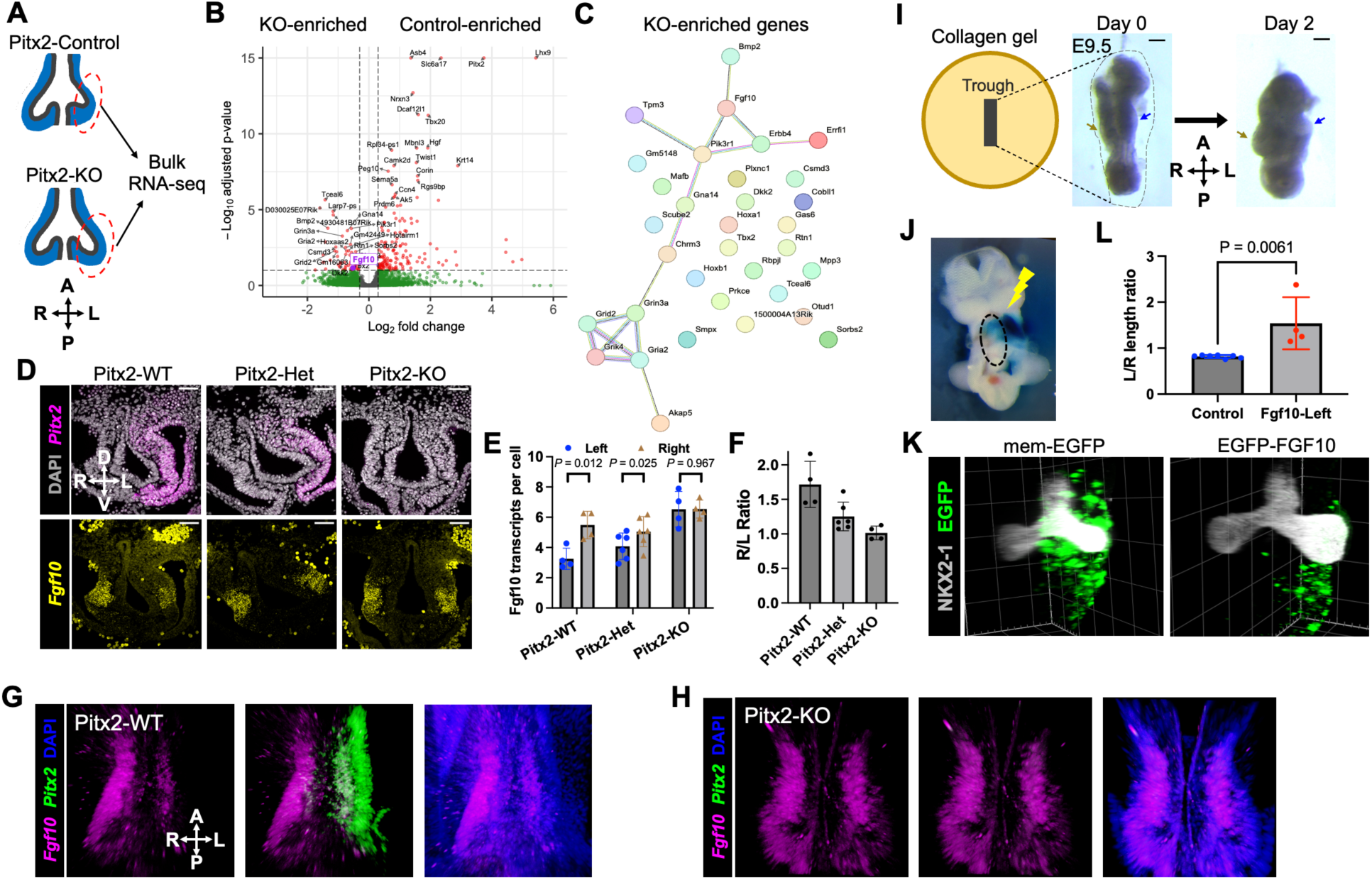
PITX2 patterns Fgf10 asymmetry in the lung bud to promote asymmetric budding. **(A)** Schematic of bulk RNA-seq of E10.5 left lung buds. **(B)** Volcano plot of differentially expressed genes between Pitx2-Control and Pitx2-KO samples (N = 4 litters). Genes with absolute log2-fold change > 0.3 and P_adj_ < 0.1 are marked red. Fgf10 is highlighted. **(C)** STRING protein-protein association network generated from genes enriched in Pitx2-KO lungs. **(D)** HCR-FISH of Pitx2 and Fgf10 in transverse sections of *Pitx2^+/+^* (WT), *Pitx2^+/-^* (Het), and *Pitx2^-/-^* (KO) at E9.5. Images are representative of N = 6 embryos from 2 litters. Scale bars: 50 µm. **(E)** Quantification of the number of Fgf10 transcripts per mesenchymal cell in the lung bud based on HCR-FISH. Each data point represents an average of 3 sections from the same embryo. N = 2 litters. P value: paired t test. **(F)** The ratio of Fgf10 transcripts per cell on the right and left side in (E). **(G,H)** Frontal views of whole mount HCR-FISH images of wild-type (G) and Pitx2-KO (H) foreguts at E9.5. Left panel: Fgf10. Middle panel: Fgf10 and Pitx2. Right panel: Fgf10 and DAPI. Images are representative of N = 4 embryos. **(I)** Ex vivo culture of embryonic lung buds in collagen gel. The blue and brown arrows indicate the left and right lung buds. Scale bars: 100 µm. **(J)** Ex vivo electroporation of the lung mesenchyme. Dyed plasmid solution was injected to the left coelom. The oval marks the lungs. **(K)** Whole mount immunofluorescence of E9.5 lung explants electroporated with membrane-tagged EGFP or EGFP-FGF10 after two days in ex vivo culture. Images are representative of N = 6 (control) or 4 (FGF10) explants. Grid size: 100 µm. **(L)** The ratio of left and right lung bud lengths quantified from (K). Each data point represents one explant. N = 2 litters. P value: unpaired t test. Error bars: standard deviation.

We next tested whether Fgf10, a critical gene in lung budding and branching^16,17^, is asymmetrically expressed in the wild-type lung, and whether this asymmetry is altered in the Pitx2-KO lung, using hybridization chain reaction fluorescence in situ hybridization (HCR-FISH) (Figures 2D, S1C and S1D). Quantification showed that Fgf10 transcript levels increased as Pitx2 dosage decreased (Figures 2E and 2F), suggesting that PITX2 reduces Fgf10 expression in the left lung bud. This early asymmetry of Fgf10 was further confirmed with whole mount HCR-FISH (Figures 2G and 2H, S1E), which is also consistent with colorimetric ISH in previous work^18–20^ and with a recent single-cell RNA sequencing dataset^21^ (Figures S1F and S1G). As Fgf10 is critical to lung budding, quantitative differences in FGF signaling intensity could potentially contribute to lung asymmetry.

To test this hypothesis, we established an explant culture system for early (E9.0-E10.0) lung buds (Figure 2I), which are challenging to culture using the conventional air-liquid interface setup^22^. Ex vivo cultured lung buds exhibit apparent normal early branches, retain high proliferation rates, and display minimal cell death as in vivo lung buds (Figures S2A-S2C). We then combined this system with ex vivo electroporation (Figure 2J), showing that overexpression of Fgf10 in the left lung mesenchyme caused the left lung bud to grow faster than the right lung, reversing the normal direction of budding asymmetry (Figures 2K and 2L). Therefore, Fgf10 asymmetry is sufficient to induce morphological asymmetry of the lung.

### Asymmetric epithelial proliferation, rather than differential directional migration, underlies early lung bud asymmetry

To determine how the left-right asymmetric FGF signaling promotes differential growth of the lung buds, we next investigated cellular processes that contribute to asymmetric lung morphogenesis. In particular, we focused on two candidate mechanisms, epithelial cell proliferation and directional migration, both of which are affected by FGF signaling later in lung development^14,18,23,24^. First, we quantified epithelial cell number in the left and right lung buds from E9.5-E12.0 and found that the asymmetry in cell number increased rapidly during E9.5-E10.5 (Figure 3A). We then performed cell cycle analysis in vivo, using a BrdU/EdU pulse-chase assay^25^, which showed that right lung epithelial cells had a significantly faster cell cycle than the left in vivo (Figures 3B and 3C). Notably, the S phase duration was not significantly different on the left versus right sides (Figure 3D), suggesting differential contribution from the G1 or G2/M phases. To directly interrogate the impact of Fgf10 on lung bud proliferation, we next returned to our ex vivo culture system (Figures 2I and 2J). Control explants exhibit a similar left-right bias in cell cycle length as observed in vivo, albeit with overall slower growth rate, due to the ex vivo conditions (Figures 3E-3G). Taking advantage of this system, we next modulated Fgf10 dosages. In explants with left-sided Fgf10 overexpression, the left lung epithelium proliferates slightly faster than the right (Figures 3H-3J), flattening the endogenous left-right difference. To evaluate the contribution of differential proliferation to lung asymmetry, we inhibited cell proliferation in the lung explant using aphidicolin. Inhibiting cell proliferation markedly reduced lung budding asymmetry (Figures 3K and 3L), suggesting that epithelial proliferation contributes substantially to the initiation of lung asymmetry.

**Figure 3.**
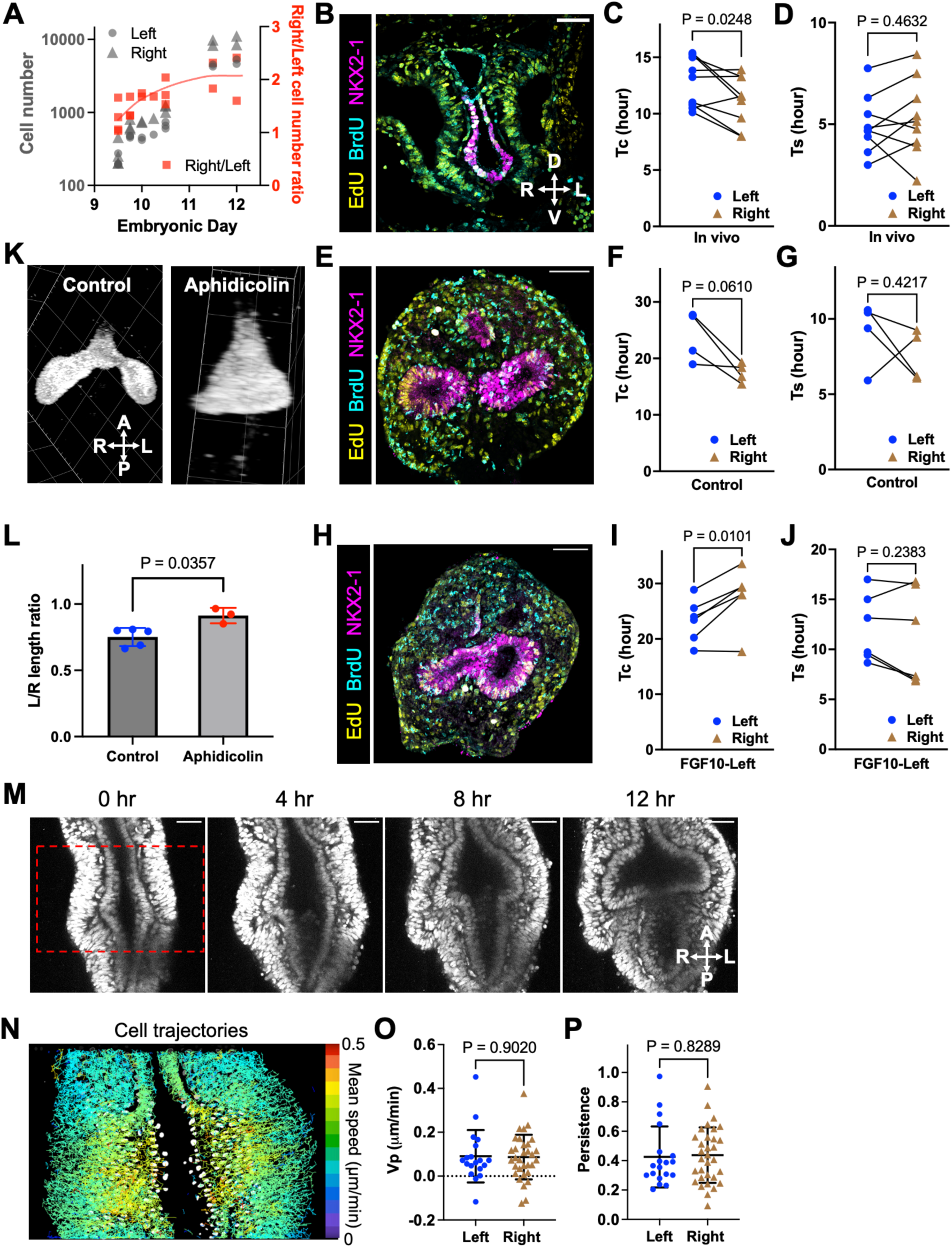
Asymmetric epithelial proliferation, rather than differential directional migration, underlies early lung bud asymmetry. **(A)** Quantification of epithelial cell numbers in the left and right lung buds and their ratio. Each data point represents one embryo. **(B,E,H)** Fluorescence image of a transverse section of an EdU/BrdU-labeled E9.5 embryo (B), a control E9.5 lung explant cultured for one day (E), or an E9.5 lung explant with left-sided Fgf10 overexpression cultured for one day (H). Image is representative of N = 3 litters. Scale bar: 50 µm. **(C,F,I)** Quantification of epithelial cell cycle length based on EdU/BrdU labeling in E9.5 embryos (C), control E9.5 lung explants cultured for one day (F), or E9.5 lung explants with left-sided Fgf10 overexpression cultured for one day (I). Each data point represents one sample. N = 3 litters. P value: paired t test. **(D,G,J)** Quantification of epithelial S phase length based on EdU/BrdU labeling in E9.5 embryos (D), control E9.5 lung explants cultured for one day (G), or E9.5 lung explants with left-sided Fgf10 overexpression cultured for one day (J). P value: paired t test. **(K)** Whole mount immunofluorescence of E9.5 lung explants treated with or without aphidicolin for one day. Images are representative of N = 3 explants. Grid size: 100 µm. **(L)** The ratio of left and right lung bud lengths quantified from (K). Each data point represents one explant. N = 2 litters. P value: unpaired t test. **(M)** Live imaging of an E9.5 lung explant in culture. Scale bar: 50 µm. **(N)** Trajectories of tracked cells within the red box in (M). Trajectories are colored by average speed. **(O,P)** Quantification of weighted velocity along the principal direction (O) and trajectory persistence (P) of tracked lung bud epithelial cells. Each data point represents one epithelial cell. P value: unpaired t test. Error bars: standard deviation.

To quantify the potential contribution of directional cell migration, we performed multiphoton live imaging of the lung explants (Figure 3M). We then used Cellpose-SAM to segment epithelial cells^26^ and tracked their movement during budding (Figures 3N, S3A-S3C, Video S2, see Methods). While the lung bud epithelium exhibited highly coherent, outward migration, the velocity of cells along the principal budding direction and the persistence of their movement were not different between the left and right sides (Figures 3O, 3P, S3D and S3E). Together, these data suggest that early lung bud asymmetry is associated with higher epithelial cell number and faster cell cycle progression on the right, which is required to maintain budding asymmetry. In contrast, directional epithelial migration occurs during budding but does not show detectable left-right difference.

### Pitx2 acts during the early budding stage to establish later branching asymmetry

Having shown that left-right asymmetry is imposed on the developing lung by Pitx2 during the early budding stage, we next asked whether the later lung branching asymmetry also requires ongoing Pitx2 activity, or whether later branching morphogenesis is affected by Pitx2 indirectly, through the changes to the tissue caused by early Pitx2 expression. Branching morphogenesis produces a stereotyped left-right asymmetric lung architecture, with fewer lobes on the left side. To test whether Pitx2 is required after the onset of branching, we generated a conditional knockout of Pitx2 (Pitx2-cKO) using the Foxg1-Cre driver^27^, which induced efficient recombination in the lung mesenchyme by E10.5. Surprisingly, although the level of PITX2 protein was significantly reduced in the *Foxg1-Cre*;*Pitx2^fl/fl^* embryos from E10.5-E11.5, lung morphology was not detectably altered at E11.5, when primary branching occurred on both sides (Figures S4A and S4B). These results suggest that continued PITX2 activity after E10.5 is not required to maintain branching asymmetry. Consistent with this interpretation, PITX2 protein level in wild-type lung mesenchyme declined over time and became barely detectable by E11.5 (Figure 4A).

**Figure 4.**
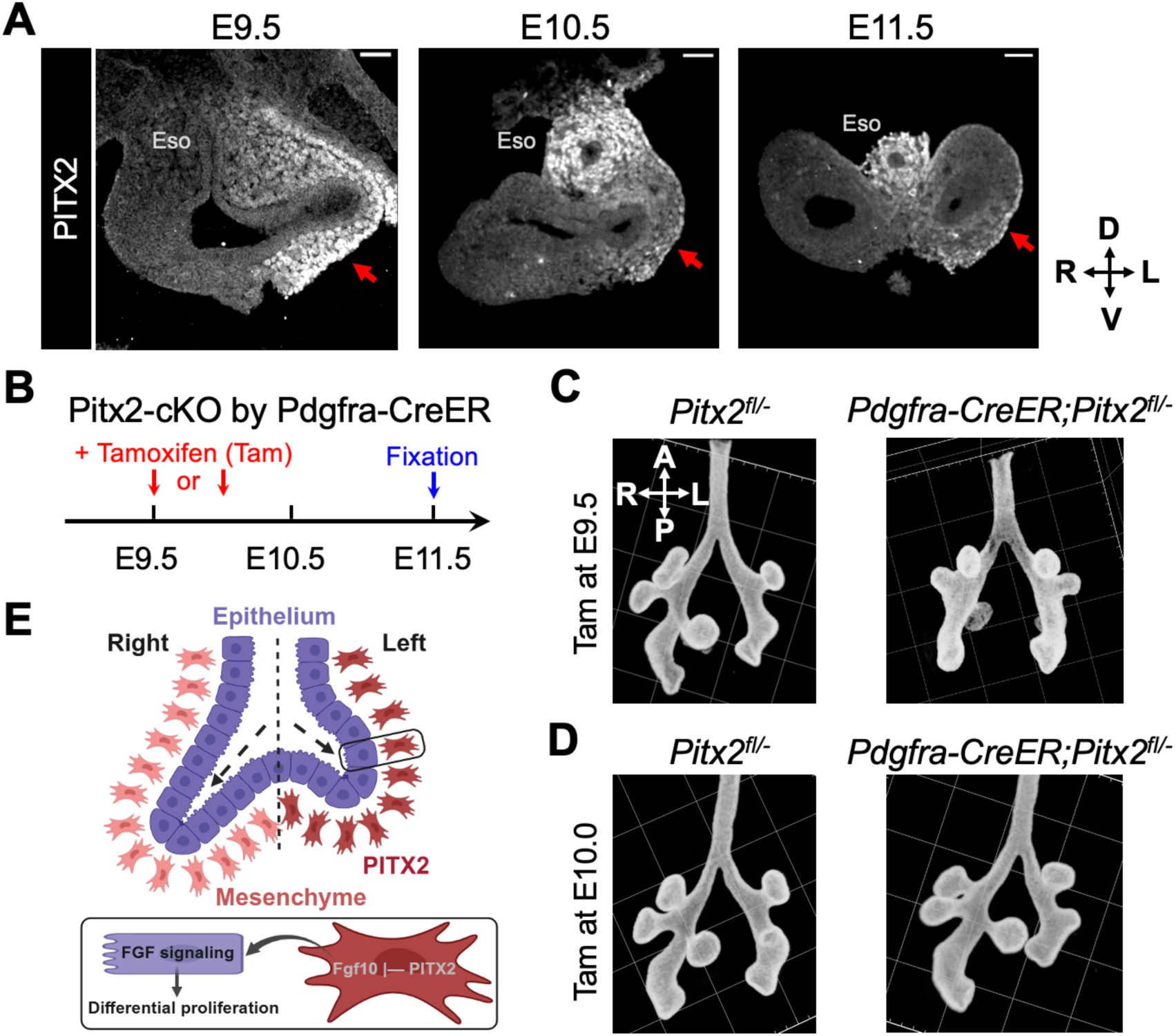
PITX2 acts during the early budding stage to establish later branching asymmetry. **(A)** Immunofluorescence images of PITX2 in transverse sections of wild-type foreguts. Arrows indicate the left lung bud. Images are representative of N = 3 embryos from independent litters. Scale bars: 50 µm. **(B)** Schematic of conditional knockout of Pitx2 by Pdgfra-CreER upon tamoxifen (tam) induction. **(C)** Whole mount immunofluorescence of E11.5 lungs of different genotypes treated with tamoxifen at E9.5. Images are representative of N = 3 litters. Grid size: 100 µm. **(D)** Whole mount immunofluorescence of E11.5 lungs of different genotypes treated with tamoxifen at E10.0. Images are representative of N = 4 litters. Grid size: 100 µm. **(E)** Model of left-right asymmetry in the early lung bud. Left mesenchyme-specific PITX2 expression inhibits Fgf10 production, which reduces FGF signaling in the left epithelium, leading to slower proliferation than the right epithelium. Schematics were created in BioRender.

To determine more precisely, when Pitx2 activity is required to establish branching asymmetry, we generated a second conditional Pitx2 knockout strain using the tamoxifen-inducible, pan-mesenchymal Pdgfra-CreER driver^28^ (Figures 4B and S4C). When a single dose of tamoxifen was administered at E9.5, the resultant Pitx2-cKO embryonic lungs showed markedly reduced branching asymmetry at E11.5, resembling the transformation of the left lung into a morphological duplication of the right lung seen in the Pitx2-KO (Figure 4C). In contrast, tamoxifen administration at E10.0 caused only a modest effect, with most Pitx2-cKO lungs retaining a wild-type-like branching pattern (Figure 4D). Therefore, the lung branching asymmetry is set by Pitx2 in the early budding stage, and subsequently, the left and right lungs undergo differential branching autonomously.

## Discussion

Our study identifies a mechanism by which left-right patterning is translated into asymmetric organ growth. During lung morphogenesis, left-specific Pitx2 expression of the mesenchyme quantitatively tunes a secreted mesenchymal signal, Fgf10, thereby biasing epithelial proliferation during the early stage of lung budding (Figure 4E).

Fgf10 has long been viewed as a central mesenchymal cue that promotes lung budding and branching across vertebrates^16–18,23,24,29^. Our findings refine this model by showing that Fgf10 is not only required for lung outgrowth, but also encodes left-right information through its dosage. Left-specific Pitx2 inhibits Fgf10 expression in the mesenchymal cells, thereby lowering the level of FGF signal seen by the left epithelium. This quantitative difference provides a mechanism for producing unequal bud elongation from an otherwise shared epithelial-mesenchymal signaling network. In other words, Pitx2 does not create a new left-specific morphogenetic mechanism but instead biases a core lung growth pathway that is already used bilaterally.

This interpretation also helps reconcile our results with previous Fgf10 rescue experiments in which ubiquitous induction of Fgf10 on an Fgf10 knockout background still generated lung asymmetry^24^. One possibility is that ubiquitously induced Fgf10 may still be inhibited by left-sided PITX2 at the transcriptional level through direct or indirect mechanisms. Elucidating these potential mechanisms will require testing of how PITX2 regulates Fgf10 in more scalable systems, e.g., in vitro differentiated lung mesenchymal cell culture^30^.

At the cellular level, our results support proliferation as the predominant cellular mechanism for the initiation of left-right asymmetry in the lungs. Although early studies on lung branching mechanisms using lung tip explants at E11.5 or later suggested that FGF signaling mainly induces cell migration instead of proliferation^14,31,32^, in vivo work showed that the effect of FGF on lung morphogenesis is highly dynamic and sensitive to developmental timing^17,33^. It is thus plausible that during the establishment of lung asymmetry at E9.5-E10.0, FGF dosage difference may be translated most strongly into cell proliferation rate, whereas during later branching, the same pathway may more prominently guide epithelial migration and branch extension.

Although lung asymmetry is ultimately manifested in differences in branching and lobation, our results indicate that the asymmetry determinant PITX2 acts surprisingly early, before stereotyped branching is morphologically apparent. PITX2 expression declines over time, and later Pitx2 deletion has little detectable effect on branching asymmetry, suggesting that PITX2 is most critical during an early competence window around lung bud initiation. Later branching can proceed with little continued PITX2 expression once an early asymmetric state has been established. We envision two possible mechanisms for how this early PITX2-dependent asymmetry is carried forward. In the ‘molecular memory’ model, transient left-sided PITX2 activity establishes a lasting mesenchymal state, potentially through establishing epigenetic changes in the genes involved in lung branching. In the ‘boundary condition’ model, PITX2 first creates a difference in bud location, size, and growth rate: and subsequent branching programs amplify this early physical asymmetry through feedback between tissue geometry and epithelial-mesenchymal interactions^10^. Future studies integrating molecular-scale dynamics with tissue geometry will be needed to answer how branching asymmetry is maintained.

## Limitations of the study

Although our data identify Fgf10 and FGF signaling as key PITX2-sensitive outputs, PITX2 likely regulates additional mesenchymal genes that may contribute to asymmetric growth or branching, as many differentially expressed genes regulated by PITX2 have yet unclear functions. The regulatory mechanism connecting PITX2 to reduced Fgf10 expression also remains unresolved. PITX2 may directly repress Fgf10 transcription or act through intermediate molecular regulators.

## Acknowledgements

We thank the MicRoN Facility of Harvard Medical School for microscopy resources, the Biopolymers Facility of Harvard Medical School for high-throughput sequencing, and the Center for Comparative Medicine of Harvard Medical School for the maintenance of mouse colonies. We thank Miram Meziane and Mackenzie Litz of the Li group, and members of the Tabin group for discussion and suggestions. We also thank Meng Zhu for the Fgf10 HCR probes and Dawei Sun for discussion of the work. This work was funded by the National Institutes of Health grant R01HD087234 to C.J.T. R.Y. acknowledges support from the Helen Hay Whitney Foundation. J.H. was funded by the Four Directions Summer Research Program at Mass General Brigham. P.L. was funded by the National Institutes of Health grant DP2HD108777 and Allen Family Philanthropies.

## Author contributions

R.Y. and C.J.T. conceived the project. R.Y. performed the experiments and data analysis. J.H. contributed to cell number and cell cycle quantification. P.L. gave advice on experimental design and data analysis. R.Y. and C.J.T. wrote the manuscript. All authors read and commented on the manuscript.

## Declaration of Competing Interests

The authors declare no competing interests.

## Methods

All animal studies were performed in compliance with the protocols approved by the Institutional Animal Care and Use Committee at Harvard Medical School.

### Mouse embryos

The following mouse strains were obtained from the Jackson Laboratory: C57BL/6J (#000664), *ROSA^nT-nG^* (nTnG, #023537)^34^, *Pitx2^tm1Rsd^* (Pitx2^+/-^, #028631)^5^, *Pitx2^tm1.1Sac^* (Pitx2^fl/+^, #018120)^3^, *Foxg1^tm1.1(cre)Ddmo^* (Foxg1-Cre, #029690)^27^, *Pdgfra^tm1.1(cre/ERT2)Blh^*(Pdgfra-CreER, #032770)^28^. Pitx2-cKO by Foxg1-Cre was generated by mating *Foxg1-Cre*;*Pitx2^fl/+^* females with *Pitx2^fl/-^* males, as the *Foxg1-Cre* allele transmits more stably in females^27^, and Pitx2-cKO by Pdgfra-CreER was generated by mating *Pdgfra-CreER*;*Pitx2^fl/fl^*with *Pitx2^fl/-^* regardless of sex. Female mice of 12-16 weeks old were used for timed pregnancy and embryo collection. The morning when a vaginal plug was observed was treated as E0.5, and the precise embryo stage was determined by limb morphology during dissection, which was not impacted by the genetic perturbations used here. For Pitx2-cKO by Pdgfra-CreER, a single dose of 20 mg/mL tamoxifen (Sigma-Aldrich, T5648) in corn oil (Sigma-Aldrich, C8267) was administered intraperitoneally to the pregnant dam at the indicated time at 1 mg/10 g body weight. All mice were housed in a specific pathogen-free (SPF) facility at Harvard Medical School under a 12-hour light/12-hour dark cycle. Animals had ad libitum access to food and water.

### Plasmids

pCAGGS-memGFP was a gift from Thomas Biederer (Addgene plasmid #115502). pCAGGS-FGF10-EGFP was made by cloning EGFP-MsFGF10 (EGFP inserted after the signal sequence of mouse Fgf10 coding sequence^35^, synthesized by Twist Biosciences) into the pCAGGS-backbone, which will be deposited to Addgene. QIAGEN Plasmid Maxi Kit was used to purify plasmids from in-house prepared DH5-alpha cells.

### Early lung explant culture

We made a narrow trough in a collagen gel pad to hold the early lung explant such that the left and right lung buds were not obstructed by the substrate surface (Figure 2I). The collagen gel was made with a solution containing 1.5 mg/mL rat tail collagen (Gibco, A1048301), 1X PBS (Invitrogen, AM9625), and sodium hydroxide (Supelco, SX0607N) in water at pH 7.5. 1 mL collagen solution was spread on a 35-mm glass bottom dish (Thermo Scientific, 150680) and incubated at 37°C for 30 minutes. The solidified gel pad was equilibrated at room temperature with 2 mL dissection medium (4% fetal bovine serum and 100X-diluted penicillin-streptomycin [Gibco, 15140-122] in DMEM with HEPES [Gibco, 21063029]), and then with 1 mL culture medium (100 ng/mL EGF [R&D Systems, 236-EG], 5% fetal bovine serum [Peak Serum, PS-FB2], and 100X-diluted penicillin-streptomycin in FluoroBrite DMEM [Gibco, A1896701]).

Fresh E9.0-E9.5 mouse embryos were collected and promptly dissected in chilled dissection medium to isolate the foregut from the pharyngeal arches to the upper stomach. The lung buds were on the ventral side of the foregut (Figure 2I). An ∼1-mm long, ∼300-µm wide trough was made in the gel pad with a fine tungsten needle (Fine Science Tools, 10130-05), and the foregut was placed in the trough with the A-P and L-R axes parallel to the gel surface. The sample was incubated in 2 mL culture medium at 37°C with 5% CO_2_ for 2-3 hours before live imaging by upright multiphoton microscopy as described below. For aphidicolin treatment, 4 µM aphidicolin (Tocris, 5736) was added to the culture medium.

### Laser ablation

Laser ablation was performed with the Leica Stellaris 8 multiphoton microscope with the Insight X3 dual beam laser and the 25X/1.0NA water immersion objective. nTnG mouse lungs were excited at 1040-nm with 9%-10% power. A video before ablation was first recorded as control. Laser ablation was then performed with 800-nm excitation at 85% power in a defined region of interest (ROI) marking the mesenchyme adjacent to the lung bud epithelium. The ROI was ablated for ∼0.1 seconds per 4-second frame for 3-10 frames, with the transmission light image being recorded during ablation to monitor the progress of ablation. Once the cells were ablated, the system was switched to the imaging mode as before ablation with a recording rate of 4 seconds per frame (Video S1). Kymographs and total displacement analyses were performed in Fiji as described^15^.

### Time-lapse imaging of the early lung explant culture

nTnG mouse foregut explants were excited at 1040-nm with 9% power on the abovementioned multiphoton microscope. A Z stack of 30-60 µm with 10-µm step size was recorded for each explant, covering the frontal planes of the early lung buds. Samples were imaged every 5-10 minutes for 12-16 hours (Video S2). The XY resolution ranged from 250 nm to 500 nm per pixel.

### Ex vivo lung bud electroporation

The plasmid mixture for electroporation was made with 4 µg/µL plasmid, 0.5% Fast Green FCF (Sigma-Aldrich, F7252), and 3% sucrose (Sigma-Aldrich, S8501) in TE buffer (Qiagen, 19086). Freshly dissected mouse embryos were placed in a rectangular well made on a 1.5% agarose gel in PBS (Gibco, 10010023) in a 10-cm dish. A mouth pipette with a thin tip, made with glass capillary (FHC Inc, 30-30-0) pulled by a micropipette puller (Sutter Instrument, P-97), was loaded with the plasmid mixture, which was injected into the space between the left lung bud surface and the left body wall (Figure 2J). The embryo was in between a pair of rod electrodes (Bulldog Bio, CUY611P7-4), with the positive electrode on the embryo’s right side. Electroporation was performed with three 40-V, 3-ms, 50-ms apart poring pulses, followed by five 8-V, 10-ms, 100-ms apart transfer pulses, all towards the positive electrode (Nepa Gene, NEPA21). The electroporated embryos were further dissected to isolate the foregut for ex vivo culture as described above.

### BrdU and EdU labeling and detection

0.15 mL of 6.25 mg/mL BrdU (5-bromo-2’-deoxyuridine, Invitrogen, B23151) was injected intraperitoneally (IP) at E9.5. After 90 minutes, 0.2 mL of 5 mg/mL EdU (5-ethynyl-2’-deoxyuridine, Invitrogen, A10044) was IP injected. The mouse was euthanized 30 minutes after EdU injection, and embryos were fixed, sectioned, and stained as described in “Immunofluorescence of tissue sections”. For E9.5 lung explants, labeling was performed at 24 hours after the start of culture. BrdU (50 µM) was labeled for 120 minutes, followed by the addition of EdU (30 µM) for 30 minutes prior to fixation. EdU staining was performed with the Click-iT EdU Cell Proliferation Kit for Imaging with Alexa Fluor 488 (Invitrogen, C10337) according to the manufacturer’s protocol.

To quantify the fraction of BrdU- and EdU-positive cells, we manually counted the total number of cells labeled by NKX2-1 and the BrdU/EdU-positive cells in the left and right lung bud epithelium. The cell fractions in 4-5 consecutive sections were averaged for each sample. The length of the S phase was inferred by 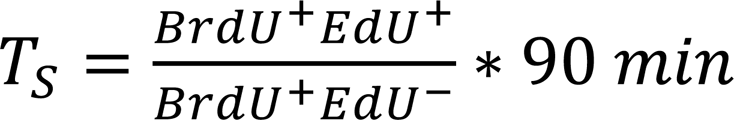 , and the length of the cell cycle was inferred by 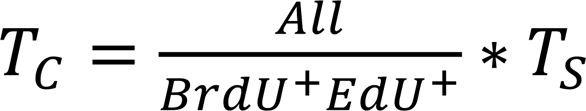 as described^25^. This calculation is valid because all lung epithelial progenitors were proliferative at this stage (Figure S2B).

### Genotyping of embryos

For Pitx2-KO and Pitx2-cKO, a mixture of genotypes was expected from a litter. We genotyped embryos by lysing the forelimb buds in 100 µL of 50 mM sodium hydroxide in water at 95°C for 30 minutes, followed by neutralization with 40 µL of 1 M Tris-HCl pH 8.0 (Invitrogen, 15568025). 1 µL of the genomic DNA preparation was added per 20 µL of reaction using the GoTaq Master Mix (Promega, M7122) according to the manufacturer’s instructions. Primers were synthesized by Integrated DNA Technologies using the genotyping sequences provided by the Jackson Laboratory. PCR reactions were analyzed by 3% agarose gel electrophoresis to determine the genotype.

### Whole mount immunofluorescence

Dissected embryonic foreguts (before E10.0) or lungs (after E10.0) were fixed in 4% paraformaldehyde (PFA, Electron Microscopy Sciences, 15710) diluted in PBS overnight at 4°C. The samples were then washed three times with PBS (10 minutes each at room temperature) and blocked with blocking buffer (10% bovine calf serum [Gibco, 16010159] and 1% Triton X-100 [Sigma-Aldrich, T8787] in PBS) for one hour at room temperature. Rabbit anti-NKX2-1 (1:300, Abcam, 76013) was diluted in the blocking buffer and labeled the samples for two days at 4°C. The samples were then washed three times with the blocking buffer (1 hour each at room temperature) and stained with donkey anti-rabbit-Alexa Fluor 647 (Jackson ImmunoResearch, 711-605-152) diluted 1:300 in the Blocking Buffer for one day at 4°C. After six washes with 5X diluted blocking buffer in PBS, the samples were serially dehydrated with 50% methanol (Fisher Scientific, A433P-4)/50% PBS, 80% methanol/20% water, and 100% methanol (1 hour each step at 4°C). Optical clearing was performed with the CytoVista Tissue Clearing kit (Invitrogen, V11322) per the manufacturer’s instruction or with dichloromethane and dibenzyl ether as described in “Whole mount hybridization chain reaction (HCR)”.

Cleared samples were transferred to a 50-mm glass bottom dish (MatTek, P50G-1.5-30-F) filled with the final clearing medium and imaged on a Nikon Ti inverted microscope with a W1 spinning disk scanner (Yokogawa, CSU-W1) using the 10x or 20x objective and 647-nm excitation. A Z-stack of ∼0.6-1.0 µm step size was acquired for each sample. Three-dimensional (3D) rendering of the image was performed in the Arivis Vision4D software (Zeiss), with the Z step size multiplied by 1.5 to correct for the refractive index mismatch.

To count cells in the left and right lung buds (Figure 3A), the 3D image was rotated such that the two lung buds can be bisected into sub-volumes. Computationally isolated lung buds were processed with Blob Finder in Arivis Vision4D to segment NKX2-1-positive nuclei with manual parameter adjustment and validation.

### Whole mount hybridization chain reaction (HCR)

Dissected embryonic lungs were fixed in 4% paraformaldehyde at room temperature for 1 hour, washed three times with PBS, and serially dehydrated to methanol as described above. For probe hybridization, the samples were treated with 70% ethanol in PBS for 1 hour at room temperature, incubated with the HCR Wash Buffer (Molecular Instruments) for 10 minutes at room temperature, and incubated with the HCR Hybridization Buffer (Molecular Instruments) for 30 minutes at 37°C. HCR probes were diluted to 40 nM in 100 µL of hybridization buffer, incubating the samples overnight at 37°C without rocking. Labeled samples were washed twice with the wash buffer, then twice with 5X SSCT (5X Saline-Sodium Citrate buffer [Invitrogen, 15557044] and 0.1% Tween 20 [Sigma-Aldrich, P9416] diluted in PBS), and finally with the HCR Amplification Buffer (Molecular Instruments), 20 minutes each at room temperature. The HCR amplifiers with Alexa Fluor 647 or Alexa Fluor 546 (Molecular Instruments) were denatured at 95°C for 90 seconds and annealed at room temperature for 30 minutes. The amplifiers were diluted in the amplification buffer at 100 nM per strand, which was incubated with the sample overnight at room temperature with rocking. For sectioning, the sample was cryo-protected and embedded in 1:1 O.C.T. Compound (Sakura Finetek, 4583):30% sucrose (Sigma-Aldrich, S8501) in PBS. For whole mount imaging, samples were washed with 5X SSCT with 10 µg/mL DAPI (Invitrogen, D1306), then with 500 mM Tris-HCl pH 7.0 (Invitrogen, AM9850G), dehydrated with methanol, and cleared with dichloromethane (Sigma-Aldrich, 270997) and dibenzyl ether (Sigma-Aldrich, 33630) according to the published protocol^36^. The sequences of HCR probes are provided in Table S2.

### Immunofluorescence of tissue sections

Whole embryos, dissected foreguts and lungs, or cultured lung explants were fixed in 4% paraformaldehyde diluted in PBS overnight at 4°C. For immunostaining of PITX2, the tissue was fixed in 2% trichloroacetic acid (TCA, Sigma-Aldrich, T6399) dissolved in Tris-buffered saline (TBS, 150 mM NaCl [Invitrogen, AM9760G] and 50 mM Tris-HCl pH 7.5 [Invitrogen, 15567027]) for 2 hours at room temperature. Fixed tissue was embedded, sectioned, and immunostained as described^15^. For staining BrdU, the sections were treated with 4 M HCl (Sigma-Aldrich, 320331) for 20 minutes at room temperature and washed with PBS before blocking. Whole mount HCR-stained samples were sectioned and stained with 10 µg/mL DAPI for 30 minutes at room temperature before mounting in the ProLong Diamond Antifade Mountant (Invitrogen, P36970).

Primary antibodies used were: rabbit anti-NKX2-1 (1:300, Abcam, 76013), sheep anti-PITX2 (1:100, R&D Systems, AF7388), chicken anti-GFP (1:500, Abcam, 13970), mouse anti-BrdU (1:100, Santa Cruz Biotechnology, sc-32323), rabbit anti-cleaved caspase 3 (1:300, Cell Signaling Technologies, 9661), rat anti-Ki-67 (1:100, Invitrogen, 740008T). Dye-conjugated secondary antibodies were from Jackson ImmunoResearch (with Alexa Fluor 647, Cy3, or Alexa Fluor 488) and used at 1:300 dilution.

### Bulk RNA sequencing (RNA-seq)

#### Library preparation and sequencing

Four litters of E10.5 *Pitx2^+/-^* X *Pitx2^+/-^*embryos were collected, and the left lung buds of *Pitx2^-/-^* vs. *Pitx2^+/+^* or *Pitx2^+/-^* were pooled from each litter as one replicate. *Pitx2^-/-^* embryos were identified by the highly penetrant heart looping defect^5^, while *Pitx2^+/+^* and *Pitx2^+/-^* embryos were phenotypically indistinguishable. Lung dissections were performed promptly as described above, and the left lung bud was cut off with a surgical blade (Aspen Surgical Products, 371111). Dissected lung buds were lysed to extract total RNA using the RNeasy Mini Kit (QIAGEN, 74104). Library construction was performed with 40 ng total RNA per sample with the HyperPrep with Ribo-Erase kit (Roche) by the Biopolymers Facility at Harvard Medical School. The 200-nM library was sequenced with a NextSeq 2000 P2 300-cycle system (Illumina) with 400 million reads at 2×150 bp.

#### Differentially expressed gene (DEG) analysis

Bulk RNA-seq count data from E10.5 left lung buds were analyzed in RStudio. The count matrix consisted of eight samples, corresponding to four Pitx2-Control and four Pitx2-KO lung bud samples. Lowly expressed genes were filtered, and differential expression analysis was performed using DESeq2^37^. Raw and normalized count distributions were inspected by boxplots, and sample relationships were assessed by principal component analysis (PCA). Differential expression results were extracted from DESeq2 and ordered by adjusted P value. Normalized counts were appended to the DESeq2 result table and exported for downstream analysis. Volcano plots were generated using EnhancedVolcano (https://github.com/kevinblighe/EnhancedVolcano), with genes considered significant using an adjusted P value cutoff of 0.1 and an absolute log2 fold-change cutoff of 0.3 (Table S1). The 20 most significant upregulated and 20 most significant downregulated genes were labeled, and Fgf10 was highlighted separately. For visualization, adjusted P values smaller than 10^-15^ were capped to limit the y-axis range.

#### STRING gene network analysis

STRING protein-protein association network was generated from genes identified in DEG analysis using default parameters^38^. Nodes represent proteins encoded by the indicated genes, and edges represent known or predicted functional associations.

### Re-analysis of published single-cell RNA-sequencing (scRNA-seq) data

E10.0 whole mouse embryo scRNA-seq dataset was obtained from Qiu et al^21^. A Seurat object was generated from raw counts, with genes detected in fewer than 20 cells and cells with fewer than 200 detected features excluded. Cells were further filtered to retain those with 300-6000 detected genes and <20% mitochondrial transcripts. Counts were log-normalized, variable genes were identified, and data were scaled before PCA, UMAP embedding, nearest-neighbor graph construction, and Leiden clustering. Splanchnic mesenchymal populations were identified using markers Pdgfra and Ptch1. Lung mesenchymal progenitors were identified by sub-clustering mesenchymal cells and scoring for Foxf1, Bmp4, Wnt2, and Isl1 expression. Within this population, cells were classified as Pitx2-positive or Pitx2-negative using a normalized Pitx2 expression threshold of 0.5.

### Cell tracking and trajectory analysis

Live imaging stacks of nTnG embryos were corrected in Fiji using Enhance Contrast, and drift-corrected by StackReg (https://bigwww.epfl.ch/thevenaz/stackreg/). Nuclear segmentation was performed in a modified Jupyter Notebook by Cellpose-SAM^26^. Segmented cell masks were shrunk by 2-fold using a custom code in MATLAB (Mathworks) to facilitate tracking. Cell tracking was performed using the TrackMate plugin^39^ in Fiji to extract the spatial coordinates of individual cells, as well as the linkage edges defining their lineage histories. The resulting spots and edges data were imported into MATLAB for downstream kinematic analysis. To specifically analyze the migration of epithelial cells, a polygonal ROI was defined in Fiji and imported to MATLAB. The principal direction of motion was determined by calculating the unit vector of the sum of all spatial displacements across all branches. The migration velocity of each lineage was then calculated by projecting the velocity vector of its constituent branches onto this principal axis, yielding a time-weighted average projected velocity (Vp). Track persistence (directionality ratio) was calculated for each branch as the ratio of the net Euclidean displacement to the total accumulated path length, and then averaged to provide a single persistence value per lineage.

### Statistics and reproducibility

Sample sizes and statistical tests are indicated in the figure legends. The sample sizes were not pre-determined. The sex of embryos was not determined as lung asymmetry is highly consistent regardless of sex. GraphPad Prism was used to plot the data.

## Data availability

The RNA-seq data of Pitx2-Control and Pitx2-KO lung buds is available at the Gene Expression Omnibus (GSE347452).

## Code availability

The custom codes used for cell tracking and trajectory analysis are available at GitHub (https://github.com/yanrui21/Cell_Kinetics_CellposeSAM_TrackMate).

## Description of supplementary files

**Video S1**. Tissue dynamics before and after laser ablation of the sub-epithelial mesenchyme in an E9.5 nTnG lung explant, related to Figures 1B-1D. Scale bar: 50 µm.

**Video S2**. Live imaging of an E9.5 nTnG lung explant in ex vivo culture with nuclear segmentation and tracking, related to Figures 3M-3P. Scale bar: 50 µm.

**Table S1**. Differentially expressed genes in between Pitx2-Control and Pitx2-KO. Genes with absolute log2-fold change > 0.3 and P_adj_ < 0.1 are included. Related to Figures 2A-2C.

**Table S2**. Sequences of the HCR probes for Fgf10, Pitx2, and Etv5.

## Figures

**Figure S1.**
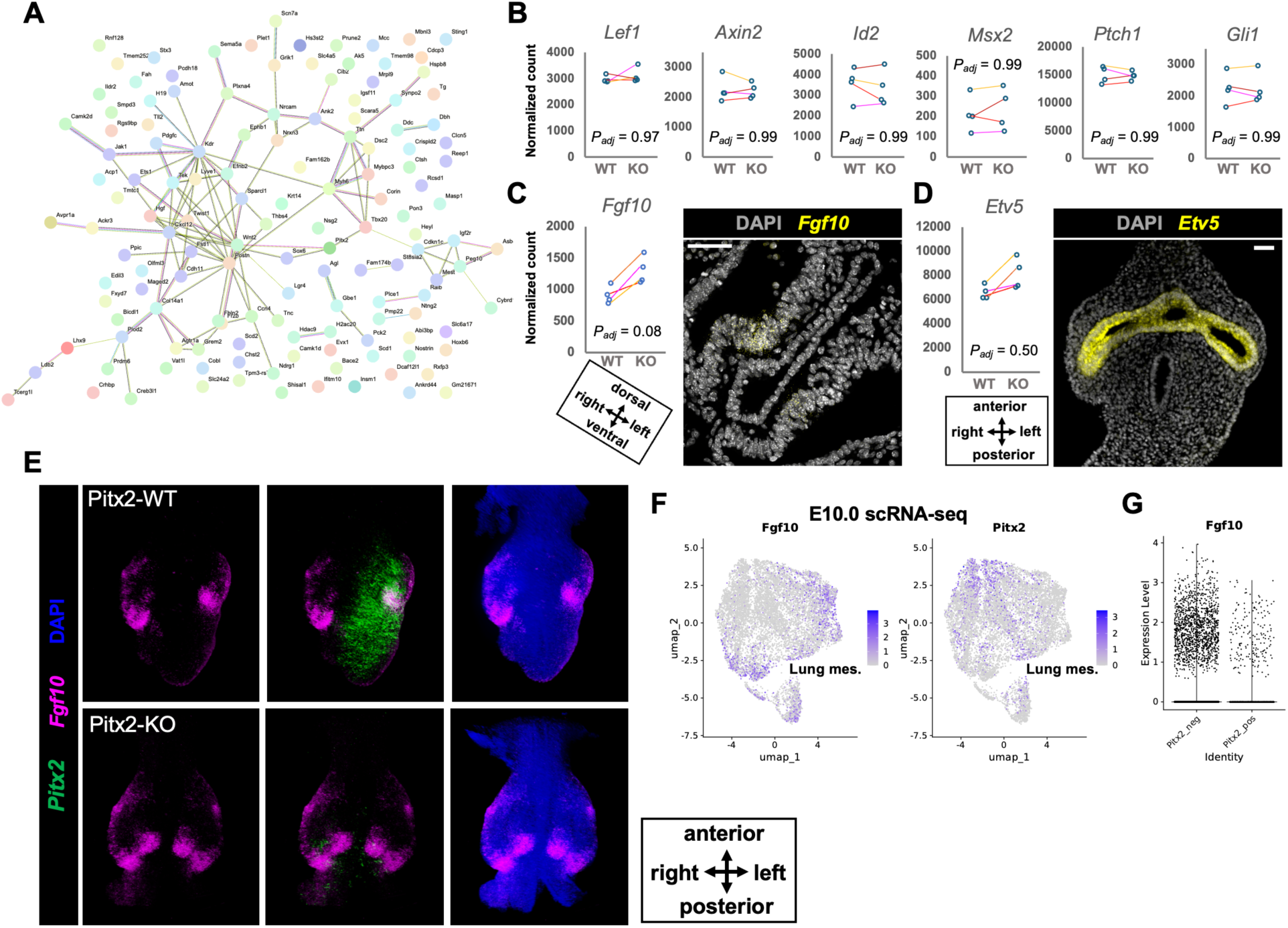
PITX2 patterns Fgf10 asymmetry in the lung bud, related to Figure 2. **(A)** STRING protein-protein association network generated from genes enriched in wild-type lungs in the RNA-seq dataset. **(B)** Quantification of WNT (Lef1, Axin2), BMP (Id2, Msx2), and SHH (Ptch1, Gli1) signaling activity based on the expression of key effector genes in the RNA-seq dataset. N = 4 litters. **(C)** Quantification of Fgf10 expression in the RNA-seq dataset and HCR-FISH of Fgf10 in a transverse section of an E9.5 embryo. Image is representative of N = 6 embryos. Scale bar: 50 µm. **(D)** Quantification of Etv5 expression in the RNA-seq dataset and HCR-FISH of Etv5 in a transverse section of an E9.5 embryo. Image is representative of N = 3 embryos. Scale bar: 50 µm. **(E)** Frontal views of whole mount HCR-FISH images of E10.5 lungs. Images are representative of N = 4 embryos. **(F)** Seurat FeaturePlots of Fgf10 and Pitx2 in the lung mesenchyme cell cluster in an E10.0 mouse embryo scRNA-seq dataset^21^. **(G)** Expression level of Fgf10 in Pitx2-positive and Pitx2-negative cells in (F).

**Figure S2.**
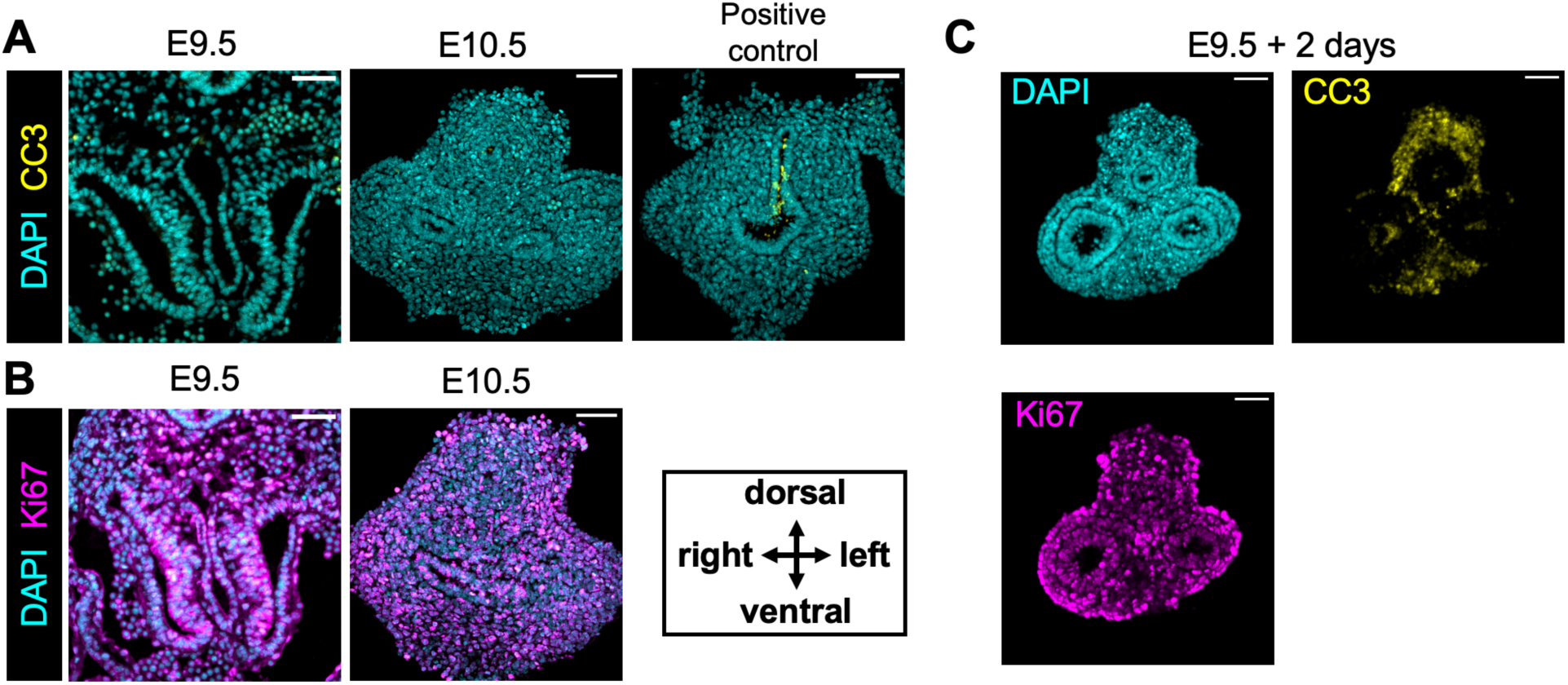
Cell proliferation and cell death in the lung bud in vivo and ex vivo, related to Figures 2 and 3. **(A)** Immunofluorescence of cleaved Caspase 3 in transverse sections of E9.5 and E10.5 embryos at the lung bud level. Positive control shows the level of tracheal-esophageal separation^15^. Images are representative of N = 3 embryos. Scale bars: 50 µm. **(B)** Immunofluorescence of Ki-67 in transverse sections of E9.5 and E10.5 embryos at the lung bud level. Images are representative of N = 3 embryos. Scale bars: 50 µm. **(C)** Immunofluorescence of cleaved Caspase 3 and Ki-67 in transverse sections of an E9.5 lung explant cultured for two days. Images are representative of N = 3 explants. Scale bars: 50 µm.

**Figure S3.**
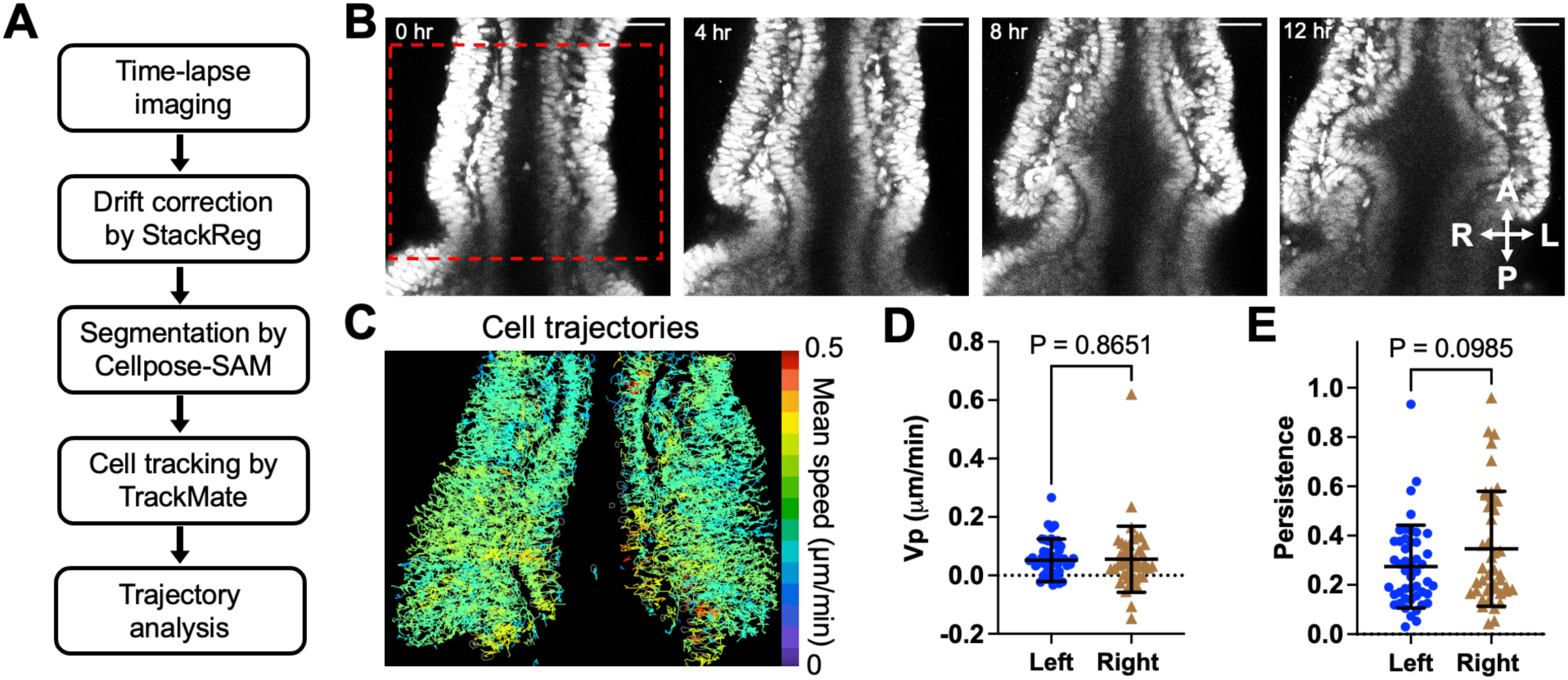
Live imaging and analysis of lung budding ex vivo, related to Figure 3. **(A)** Workflow of live imaging analysis. **(B)** Additional example of live imaging of an E9.5 lung explant in culture. Scale bar: 50 µm. **(C)** Trajectories of tracked cells within the red box in (B). Trajectories are colored by average speed. **(D)** Quantification of weighted velocity along the principal direction of tracked lung bud epithelial cells. P value: unpaired t test. **(E)** Quantification of trajectory persistence of tracked lung bud epithelial cells. P value: unpaired t test. Error bars: standard deviation.

**Figure S4.**
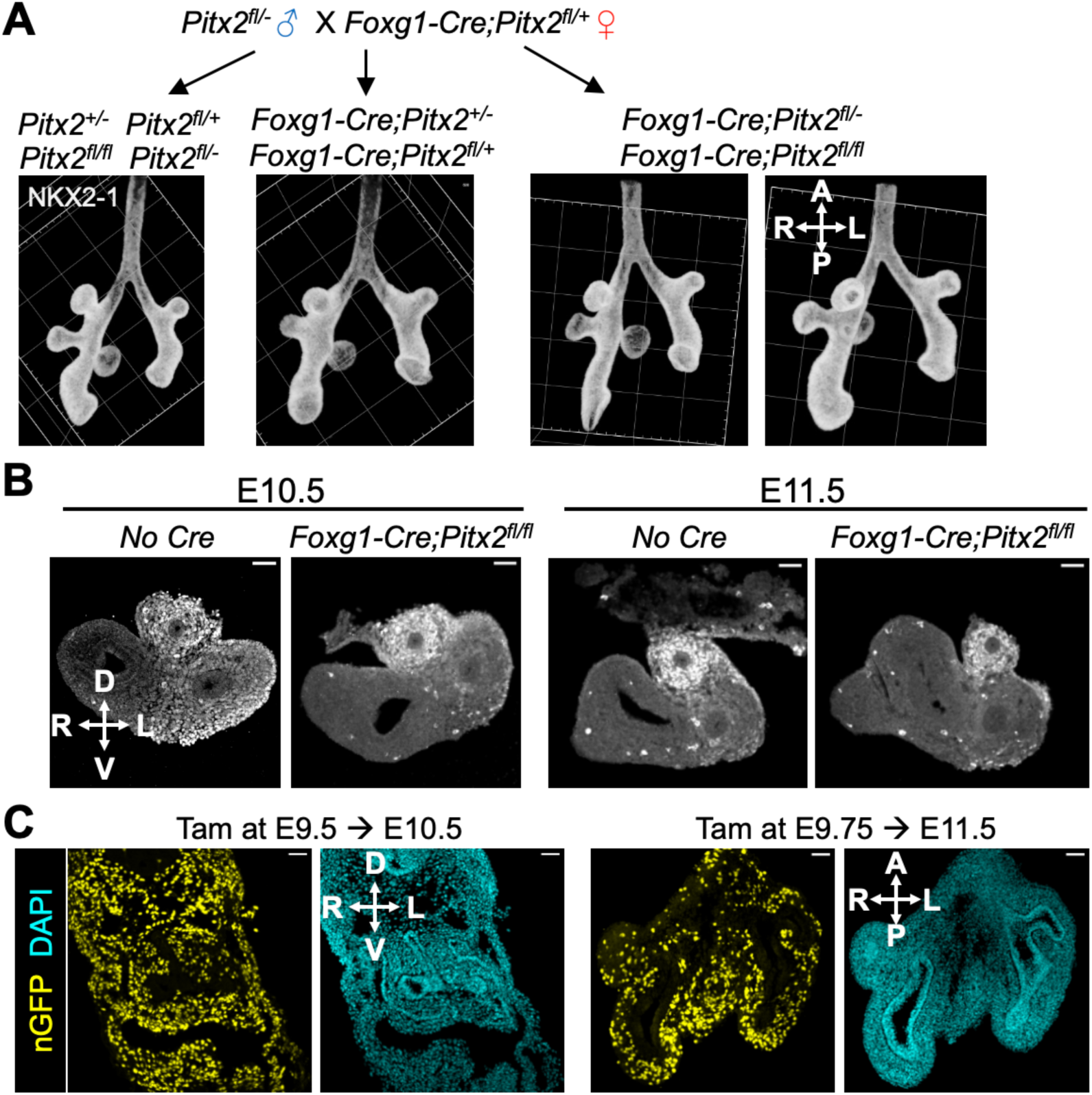
Pitx2 depletion after E10.5 does not alter lung asymmetry, related to Figure 4. **(A)** Whole mount immunofluorescence of E11.5 Foxg1-Cre-mediated Pitx2-cKO lungs of different genotypes. Images are representative of N = 3 litters. Grid size: 100 µm. **(B)** Immunofluorescence of PITX2 in transverse sections of Foxg1-Cre Pitx2-cKO lungs of different genotypes. Images are representative of N = 4 litters. Scale bars: 50 µm. **(C)** Immunofluorescence of transverse sections (left) and frontal sections (right) of *Pdgfra-CreER*;*nTnG/+* embryos treated with tamoxifen. Images are representative of N = 3 litters. Scale bars: 50 µm.

## Notes

### Competing Interest Statement

The authors have declared no competing interest.

### Summary of Updates

Figures updated. GEO accession number provided.

